# Seizure-like behavior and hyperactivity in *napb* knockout zebrafish as a model for autism and epilepsy

**DOI:** 10.1101/2025.03.24.644881

**Authors:** Kyung Chul Shin, Waseem Hasan, Gowher Ali, Doua Abdelrahman, Tala Abuarja, Lawrence W Stanton, Sahar I. Da’as, Yongsoo Park

## Abstract

We identified *N*-ethylmaleimide-sensitive factor attachment protein beta (*NAPB*) as a potential risk gene for autism and epilepsy. Notably, Qatari monozygotic triplets with loss of function mutations in *NAPB* exhibit early onset epileptic encephalopathy and varying degrees of autism. In this study, we generated NAPB zebrafish model using CRISPR-Cas9-sgRNAs technology for gene editing of the two orthologs *napba* and *napbb*. We observed that *napb* crispants (CR) show shorter motor neuron axons length together with altered locomotion behavior, including significant increases in larvae total distance traveled, swimming velocity, and rotation frequency, indicating that these behavioral changes effectively mimic the human epileptic phenotype. We applied microelectrode array (MEA) technology to monitor neural activity and hyperexcitability in the zebrafish model. The *napb* CR shows hyperexcitability in the brain region. By combining behavioral tests with electrophysiological MEA assays, the established NAPB zebrafish model can be employed to study the pathophysiological mechanisms of ASD and epilepsy to screen potential therapeutic drugs.

## Introduction

Autism spectrum disorder (ASD) is a complex neurodevelopmental syndrome defined by repetitive behaviors and communication impairments^1^. Comorbidities are common in ASD, including epilepsy^2^. Our research has identified *N*-ethylmaleimide-sensitive factor attachment protein beta (*NAPB*) as a potential risk gene for ASD^3,4^. Homozygous autosomal genetic variants in *NAPB*, inherited from heterozygous parents, were found in ASD triplets from the Qatari population. These affected male monozygotic triplets also exhibited early-onset epileptic encephalopathy with varying severity and pharmacological repsonses^3,4^. Proband-specific, parent-specific, and genetically corrected *NAPB* pluripotent stem cell lines were generated^5^. Human neurons derived from these lines showed loss of NAPB function in neurons from the probands^4^. Studies by two independent international groups have also identified two additional unique loss-of-function mutations in *NAPB* associated with epilepsy^6,7^, further supporting the role of *NAPB* mutations in ASD and epilepsy.

NAP, also known as soluble NSF attachment protein (SNAP), is an essential cofactor of NSF ATPase, facilitating the disassembly of the SNARE complex and thereby increasing the availability of free SNARE proteins for subsequent fusion reactions^8^. NAP-alpha (NAPA) and NAP-beta (NAPB) share 83% similarity, with molecular weights of 33 kDa and 34 kDa, respectively^9^. NAPB is specifically expressed in the brain, while NAPA is found in a wide range of tissues^9,10^. Correlating with the presence of epilepsy in patients with *NAPB* mutations, *Napb* knockout (KO) mice exhibit severe recurrent epileptic seizures postnatally^10^. However, *NAPB* deletion does not affect synaptic transmission and short-term plasticity at the cellular level^10^. Similarly, we have observed no defects in neuronal activity and calcium influx in *NAPB* mutant human cortical neurons generated from pluripotent stem cell lines^4^, suggesting that cellular models may not sufficiently recapitulate the epileptic phenotype. Developing animal models of *NAPB* genetic variants remains challenging for studying the pathophysiological mechanisms of ASD and epilepsy.

Zebrafish (*Danio rerio*) are commonly used to study epilepsy and neurodevelopmental disorders due to their genetic similarity with humans^11,12^. Genes associated with these conditions are conserved between zebrafish and humans^13,14^. Moreover, zebrafish exhibit quantifiable behaviors that can model seizures, hyperactivity, and other neurodevelopmental symptoms^11,15^. The optical transparency and well-defined central nervous system of zebrafish further facilitate modeling seizures and epilepsy^11,12^. In this study, we generated NAPB zebrafish model using CRISPR-Cas9 gene manipulation targeting the two zebrafish orthologs *napba* and *napbb* to investigate the pathology of NAPB-associated epilepsy.

A microelectrode array (MEA) is a functional and electrophysiological assay that measures extracellular field potentials generated by neural activity^16^. Multiple electrodes independently measure across the central nervous system, enabling the evaluation of neural networks in a non-invasive and label-free manner^16^. MEA is a valuable tool for monitoring neural activity in zebrafish *in vivo*^17–19^, allowing us to study hyperactivity and hyperexcitability in zebrafish models of epilepsy.

Here, we generated *napb* CR as an animal model to investigate neurological presentations, including seizure-like phenotypes and epilepsy-related to *NAPB* genetic variation. We utilized MEA technology to monitor neural hyperexcitability in *napb* CR, combining behavioral tests and electrophysiological functional assays to study the pathophysiology of epilepsy and related neurological disorders.

## Material and Methods

### 1.1 Zebrafish care and maintenance

Zebrafish (*Danio rerio*) wild-type line (AB strain) and transgenic reporter line (Tg:olig2:dsRed) adults were maintained in a recirculating aquaculture system under standard environmental conditions (the source of zebrafish is from Research Department, Sidra Medicine, Qatar): temperature at 27 °C, conductivity at 800-1000 µs and pH at 7.5 with 14 h light and 10 h dark cycle. All protocols used in these studies were approved by the local Animal Care and Use Committee and conform to the Zebrafish Policy published by the Qatar Ministry of Public Health that follows the Guide for the Care and Use of Laboratory Animals published by the National Institutes of Health. Experiments performed on zebrafish were approved by the IACUC Office of Sidra Medicine, approval SIDRA-2023-001 and SIDRA-EXEMPT-2024-002. All experiments were performed in accordance with the ARRIVE guidelines. For zebrafish injection experiments, embryos were collected for microinjection at the 1-cell stage and raised in 28 °C incubators. For behavioral assays, embryos were kept in E3 media (5 mM NaCl, 0.17 mM KCl, 0.33 mM CaCl_2_, 0.33 mM MgSO_4_)^20^. For Imaging assays, Tg:olig2:dsRed embryos were transferred to 75 µM N-phenylthiourea (PTU) (Sigma, P7629) media at 22 hours post-fertilization (hpf) to inhibit melanogenesis^21^. Larvae at 120 hpf were used for phenotypic examination and imaging for morphological analysis. At the end of the experiments, zebrafish larvae were euthanized by the administration of Tricaine MS-222 (168 mg/L, Western Chemical, 1029D11), followed by chilling on ice. Upon euthanasia, zebrafish carcasses were disposed of as biological waste.

### 1.2 Design of guide RNA and zebrafish microinjection

CRISPR-Cas9 gene editing was used to generate zebrafish models of *napb* knockout (KO). Zebrafish orthologs, *napba* (ENSDARG00000013669) and *napbb* (ENSDARG00000069101), were targeted using mix injections of sgRNAs. The sgRNA sequences were: MIX 1 targeting *napba*: sgRNA sequence + PAM (5’-3’): CACGCAATCATGGACAATTC CGG and sgRNA sequence + PAM (5’-3’): CTTGAACATGTTGGCAGCTC TGG; MIX 2 targeting *napbb*: sgRNA sequence + PAM (5’-3’): TTTGAACATGTTGGCGGCTC TGG and sgRNA sequence + PAM (5’-3’): GATGTGATGCTTTGCTGCGA TGG. Combined sgRNAs with Cas9 RNP mixes were injected with fluorescein or rhodamine fluorescent dye at one-cell stage embryos. At 24 hours post-fertilization (hpf), the injected embryos were screened for fluorescence. The extracted genomic DNA of pooled embryos (n=25) was analyzed using High-Resolution Melting Analysis (HRM) to confirm the early CRISPR-induced indels in the crispants using designed primers (**Supplementary Table 1**).

### 1.3 Zebrafish motor neuron length measurements

Phenotypic analysis and live imaging were conducted on the transgenic reporter line Tg:olig2-dsRed, which labels elongated spinal motor neurons at 120 hpf. Larvae were immobilized in 0.8% low melting agarose (Sigma, A9414) for imaging using Vast BioImager microscopy (Union Biometrica, Spain), as previously described^22^. The motor axon length was traced and measured for 7 axons per larva, axon numbers 7-15, using DanioScope software (Noldus, Netherlands).

### 1.4 Zebrafish neuromuscular synaptogenesis

For neuromuscular synaptogenesis, embryos were anesthetized and fixed in 4% paraformaldehyde in phosphate-buffered saline (PBS); after fixation and rinsing in PBS, embryos were incubated in 10 μg/ml fluorescently Alexa Fluor 594 conjugated with α-bungarotoxin (αBTX) to detect nicotinic acetylcholine receptors (nAChRs) located on the post-synaptic membrane at the neuromuscular junction (NMJ). αBTX (Thermofisher, Cat# B13423) diluted in blocking solution (2% BSA, 0.5% Triton X-100 in PBS). After rinsing, embryos were incubated overnight at 4 °C with primary antibody diluted in blocking solution: antibody against synaptic vesicle glycoprotein 2A (SV2) was used to detect pre-synaptic area (1:50–1:100 dilution, Developmental Studies Hybridoma Bank (DSHB), Cat # SV2)^23^.

### 1.5 Pentylenetetrazol (PTZ) treatment

The convulsant agent PTZ (P6500, Sigma-Aldrich) was prepared by dissolving it in deionized water, while the final concentration used (15 mM) was obtained by diluting the stock in Egg water. After PTZ exposure for 30 minutes in dark condition, locomotion behavior and neuronal activity were examined as described below.

### 1.6 Zebrafish locomotor behavior measurements

Zebrafish larvae locomotor activity was monitored at 120 hpf using an automated Video-Track system (Noldus, Ethovision XT, Netherlands); the behavior was assessed by measuring swimming speed, distances swam, time spent mobile and immobile, erratic movements, turn angle, etc. as described previously^24^. Light-to-dark and dark-to-light transitions are well-documented paradigms used in zebrafish epilepsy models to elicit seizure-like behaviors and increased locomotor activity. Sudden changes in lighting conditions provoke distinct behavioral responses, serving as reliable indicators of neural hyperactivity, a hallmark of epilepsy^25–27^. Larvae were placed individually in a 24-well plate in E3 medium. Larval swimming behavior was monitored in response to dark-to-light transitions after 5-min acclimatization time in the dark (12-min recoding with 2-min dark-light cycles). The Ethovision system was equipped with a BASLER camera and set to 30 frames per sec. In each cycle, we recorded the frequency of movements, distance traveled, velocity, and mobility. Larvae were incubated in 15 mM PTZ for 30 minutes (min) under dark conditions. The effects of PTZ were monitored during the 30-min incubation period, followed by a 12-min recording of dark-to-light transitions, consisting of 2-min dark-light cycles. This protocol was designed to evaluate seizure-like locomotor responses during PTZ exposure. Four technical replicates for each group were performed. The locomotor behavior was monitored, and the collected data were analyzed using Ethovision XT software.

### 1.7 Neural activity measurement using Microelectrode Array (MEA)

Zebrafish larvae were immobilized using 1.5% low melt agarose (Sigma, A9414) on 6-well CytoView MEA plates (Axion Biosystems). The agarose was dissolved in E3 Media. Once the agarose cooled but before solidifying (within 45 sec to 1 min), zebrafish larvae were quickly placed into the agarose solution and then transferred into the wells, which were filled with PBS^17–19^. Spontaneous neural activity was monitored for 10 min, followed by 30 min in the presence of 15 mM PTZ.

Extracellular recordings were conducted using Axion’s Maestro Pro platform in combination with Axion 6-well CytoView MEA plates. Each well contained an 8×8 (64-channel) electrode array with seven reference electrodes. Extracellular voltage recordings were collected at a sampling rate of 12.5 kHz per channel. A band-pass filter (200 Hz to 3 kHz cut-off frequencies) was applied, and a variable threshold spike detector was set at ±5.5 standard deviations of the root mean square (RMS) of the background noise.

MEA data for 5 minutes of recording showing active neural responses were selected for analysis. The analysis was performed using the Axion Biosystems Neural Metric Tool^28^. An electrode was considered active with a threshold of 5 spikes/min. The weighted mean firing rate was defined as the average firing rate of only the active electrodes. A burst was defined as at least 5 consecutive spikes from a single active electrode with inter-spike intervals (ISI) of less than 100 ms. Network bursts were defined as at least 5 consecutive spikes across multiple electrodes with ISIs of less than 100 ms and a minimum involvement of 10% of the electrodes.

### 1.8 PCR and qPCR

Total RNA was extracted from zebrafish larvae using the Direct-zol RNA Extraction Kit (Zymo Research), following the ‘manufacturer’s protocol. Complementary DNA (cDNA) was synthesized from 1 µg of RNA using reverse transcriptase, the RevertAid First Strand cDNA Synthesis Kit (Thermo Fisher Scientific). Gene expression for the transcripts listed in **Supplementary Table 2** was validated using conventional PCR with the PCR Master Mix (Thermo Fisher Scientific). Quantitative PCR (qPCR) was performed using the SYBR Green PCR Master Mix (Applied Biosystems), and amplification was detected using the QuantStudio 7 System (Applied Biosystems). Gene expression levels were normalized to *actb1*.

### 1.9 Statistical analysis

Data analysis was performed using OriginPro 2019 software (OriginLab Corporation, Northampton, MA, USA) and GraphPad Prism 9 (GraphPad Software, San Diego, CA, USA). Data are presented as means ± standard deviation (SD). Welch and Brown-Forsythe’s one-way ANOVA test was used to determine any statistically significant differences between three or more independent groups. A two-way ANOVA with post-hoc Tukey’s test was used to analyze the effects of two independent categorical variables, i.e., mutant and PTZ. A Repeat measure one-way ANOVA was used to compare paired groups before and after PTZ treatment, as shown in **Figure 3**. The unpaired two-tailed *t*-test was used to estimate the statistical significance between the two groups. Probabilities of *p* < 0.05 were considered significant. Outliers are removed using the Robust regression and Outlier removal (ROUT) method with Q = 1%.

## Results

### Zebrafish animal models to study *NAPB* mutations for epilepsy

Using whole exome sequencing on more than 100 ASD families in Qatar, our previous report identified *N*-ethylmaleimide-sensitive factor attachment proteins beta (*NAPB*) as a potential risk gene for autism spectrum disorder (ASD) and other neurodevelopmental disorders^4^. Homozygous recessive inheritance of genetic variants in an RNA splice site of the *NAPB* gene in triplets leads to exon-skipping and a frameshift mutation that causes a loss-of-function (LoF) in *NAPB*^3,4^. Additional NAPB LoF mutations were also linked to epilepsy^6,7^. Our observations indicate that all three probands exhibit epilepsy with varying degrees of severity and frequency, underscoring the role of *NAPB* mutations in both ASD and epilepsy, suggesting a potential genetic overlap between these conditions^3,4^.

Animal models are essential for investigating the pathophysiology of neurodevelopmental disorders and epilepsy caused by *NAPB* mutations. Here, we utilized the zebrafish model to replicate the epileptic phenotype associated with *NAPB* mutations. Zebrafish are a powerful model organism for studying epilepsy and neurodevelopmental disorders^11,12^. The human *NAPB* Gene (ENSG00000125814) has two zebrafish orthologs; i.e., *napba* (ENSDARG00000013669) with 85.9% identity (96.6% similar) in 298 aa overlap and *napbb* (ENSDARG00000069101) with 87.6% identity (97.7% similar) in 298 aa overlap (**Supplementary Figure 1A**).

Zebrafish crispants (CR) were created to knockout (KO) the two *napb* orthologs using CRISPR gene editing by injecting 1-cell stage embryos and then performing a high-resolution melting curve assay of 25-pooled embryos (**Supplementary Figure 1B-D**). Measuring motor neuron length provides critical insights into the structural and functional integrity of motor circuits, which are often disrupted in neurodevelopmental disorders and epilepsy^29^. We performed live imaging for the spinal motor neuron development and observed that the *napb* CR zebrafish model showed shorter axon length of motor neurons extending from the spinal cord and innervating the muscles when compared to the control group (**Figure 1A,B**), showing the defects on neural development and connectivity in neurodevelopmental disorders and epilepsy.

**Figure 1.**
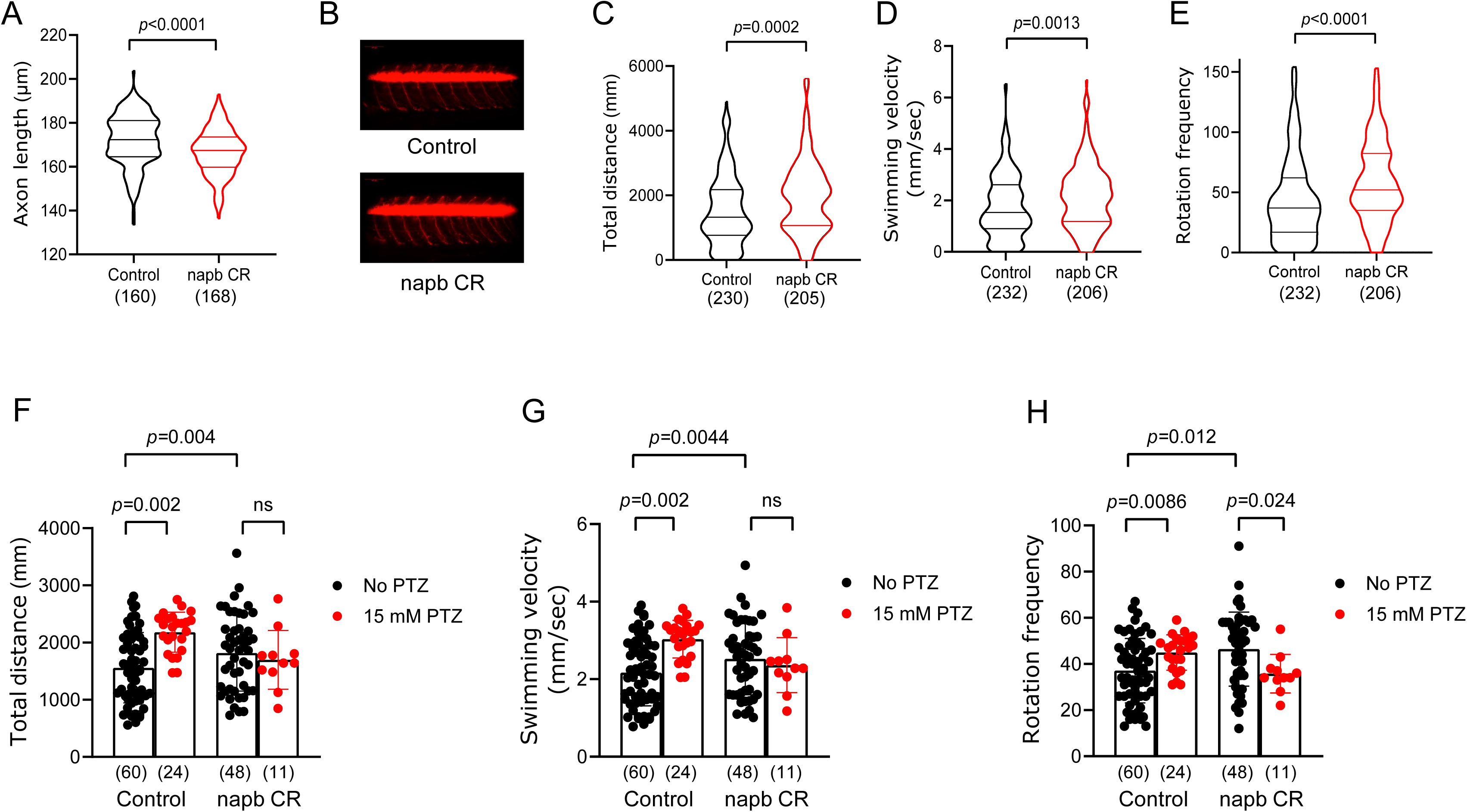
Hyperactivity and seizure-like phenotype of *napb* CR. (**A**) Axon length measured for 7 axons per embryo. (**B**) Representative lateral zebrafish larva images of control and *napb* knockout (KO) crispants (CR). (**C**) Total distance and (**D**) velocity of larvae were calculated using Ethovision software (Noldus Technologies). (**E**) Rotation frequency of larvae, defined as the number of clockwise and/or counterclockwise rotations with a threshold of 45° and a minimum distance of 1 mm. For panels **A-E**, violin plots display the median and quartiles. The number of larvae tested is shown in parentheses, and statistical analysis was performed using an unpaired two-tailed *t*-test. Outliers were excluded in (**A**). (**F-H**) PTZ induces seizures in control zebrafish larvae: (**F**) Total distance, (**G**) swimming velocity, and (**H**) rotation frequency of control larvae in the presence or absence of 15 mM PTZ. Data (**F-H**) are means ± SD. The number of larvae tested is shown in parentheses. Two-way ANOVA test with post-hoc Tukey was applied to analyze the effects of two independent categorical variables, i.e., *napb* CR and PTZ (**F, G**). Welch and Brown-Forsythe one-way ANOVA test was used to determine statistically significant differences between three or more independent groups (**H**).

Furthermore, we analyzed neuromuscular junctions (NMJs), which are key points for synaptic connectivity changes linked to hyperexcitability or seizure activity^29^. The neuromuscular synapses were examined using labeling with Synaptic vesicle glycoprotein 2 (SV2, a pre-synaptic vesicle glycoprotein essential for neurotransmitter release) and α-Bungarotoxin (αBTX), which specifically binds to post-synaptic acetylcholine receptors (AChRs). The *napb* CR showed no difference when compared to the control group. Both pre-and post-synaptic terminals demonstrated patterned clusters of axon branching within muscle layers (**Supplementary Figure 2**).

In zebrafish models of epilepsy, behavioral changes and hyperactivity can be quantified through various measures, such as alterations in locomotor behavior, including total distance traveled, swimming velocity, and rotation frequency^11,15^. The *napb* CR showed significant increases in total distance traveled (**Figure 1C**), swimming velocity (**Figure 1D**), and rotation frequency (**Figure 1E**), supporting that behavioral changes in *NAPB* zebrafish model mimic the human epileptic phenotype and highlighting that *napb* CR can be further employed to investigate the pathophysiology in ASD-related epilepsy.

We assessed the individual targeting of each ortholog, ***napba*** and ***napbb*** (Supplementary Figure 3). Locomotion analysis, including total distance traveled, swimming velocity, and rotation frequency, showed no significant effects when each ortholog was targeted separately. This finding suggests that these two orthologs may exhibit compensatory effects, where the remaining gene compensates for the loss of function when either ***napba*** or ***napbb*** is mutated individually, thereby masking any potential phenotypic effects. However, when both genes are mutated simultaneously in ***napb*** crispants, the defects become more pronounced. This concept of functional redundancy and compensation among duplicated genes is supported by previous zebrafish studies^30^. Based on this, we generated a zebrafish ***napb*** CR by simultaneously targeting both ***napb*** orthologs through the direct delivery of a ribonucleoprotein (RNP) complex consisting of Cas9 protein and single guide RNA (sgRNA) targeting ***napba*** and ***napbb*** (Supplementary Figure 1).

### PTZ-induced seizure activity in control zebrafish, but not napb CR

Pentylenetetrazol (PTZ) is a chemical compound that induces seizures by antagonizing the activity of GABA type A receptors, thereby reducing the inhibitory effect of GABA and leading to increased neuronal excitability and seizure activity in zebrafish^31,32^. Hyperactivity and seizure-like behaviors are common responses to PTZ, a seizure-inducing agent^31^.

The gene expression of various GABA-A receptor subunits in zebrafish larvae was confirmed using reverse transcription quantitative PCR (qPCR). (**Supplementary Figure 4A**). As expected, the PTZ treatment (30 min incubation with 15 mM PTZ) caused hyperactivity and seizure-like behavior in control zebrafish, evidenced by significantly increased total distance traveled (**Figure 1F**), swimming velocity (**Figure 1G**), and rotation frequency (**Figure 1H**). The zebrafish behavior was assessed over a 12-min period immediately following a 30-min PTZ incubation. Detailed quantification, visualized in 2-min time bins, revealed consistent behavioral patterns throughout the 12-min total recording. The *napb* CR exhibited significantly higher total distance traveled, velocity, and rotation over time compared to the control group prior to PTZ exposure (**Supplementary Figure 5A-C**). Following PTZ exposure, the control group demonstrated a pronounced response, characterized by a significant increase in total distance traveled, velocity, and rotation (**Supplementary Figure 5**).

Visualization of locomotor behavior using red tracking lines over time during dark-light cycles revealed distinct swimming patterns for the *napb* CR zebrafish before and after PTZ incubation. Control larvae displayed organized swimming along the well periphery, whereas *napb* CR zebrafish demonstrated irregular swimming within the well with increased rotations (**Supplementary Video 1**).

To further evaluate the behavior of *napb* CR zebrafish in the presence of PTZ, seizure-like behavior was analyzed in 2-min intervals immediately following 15 mM PTZ incubation (**Supplementary Figure 6**). During the first 8 minutes of PTZ exposure, *napb* CR zebrafish showed significantly higher total distance traveled and swimming velocity, as well as a significantly increased rotation frequency, compared to the control group’s locomotor behavior. However, the effect of PTZ diminished over time, possibly due to the occurrence of catatonic periods. (**Supplementary Figure 6**).

### Neural hyperactivity and hyperexcitability in *napb* CR using MEA

Next, we investigated the neural activity and network formation in zebrafish using MEA technology. The MEA consists of a grid of 64 tightly spaced electrodes that detect extracellular field potentials generated by the neural activity of zebrafish (**Figure 2**). MEA enables simultaneous monitoring of neural activity from different locations across the entire body of the zebrafish, allowing for the detection of propagation and synchronization of neural activity, measuring both neuronal activity and neural network formation^17–19^.

**Figure 2.**
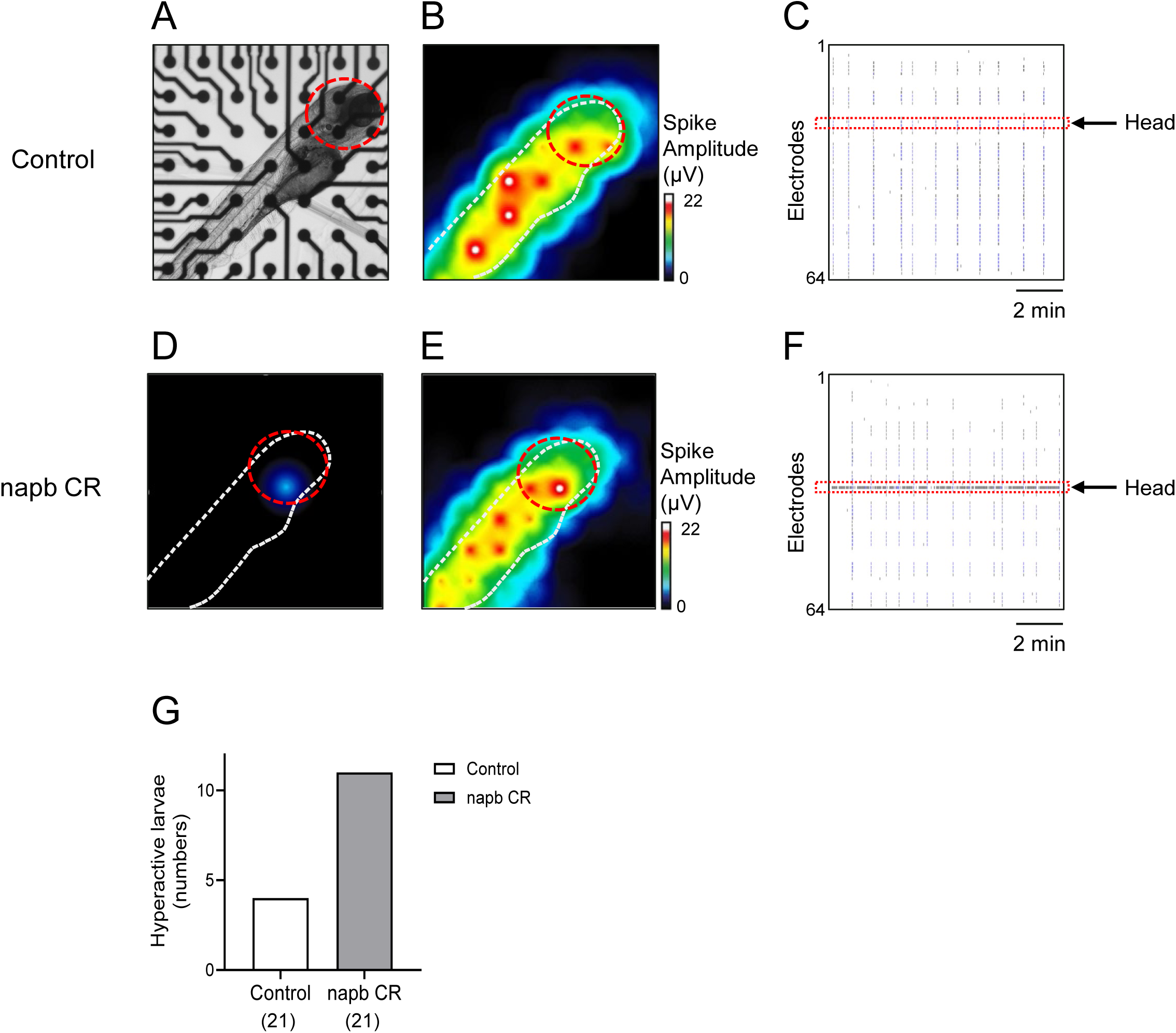
Electrical hyperexcitability in *napb* CR. Neural activity and neural network were monitored using MEA. (**A**) Inverted microscope image of a control zebrafish larva, laterally immobilized on the electrodes of MEA plates. (**B**) Heat map of spontaneous spike amplitude (µV) of a control zebrafish, showing that detected extracellular activity aligns with the zebrafish’s orientation (white dot). The red circle indicates the head region. (**C**) Well-wide raster plot illustrating synchronized spinal cord and brain neural activity in the control zebrafish. The red dotted line indicates neural activity in the brain region. (**D-F**) Hyperactive neural activity of *napb* CR. (**D, E**) Heat map of spontaneous spike amplitude (µV) and (**F**) raster plot of *napb* CR. (**G**) The number of zebrafish larvae exhibiting hyperactive neural activity in the brain region; control (n = 21) and *napb* CR (n = 21).

Zebrafish were laterally immobilized with low melting agarose on the surface of the MEA electrodes (**Figure 2A**), which recorded the spontaneous neural activity of the zebrafish. MEA electrodes detect changes in extracellular local field potentials evoked by membrane depolarization and activation of the nervous system^16^. A heat map indicated that spontaneous neural activity aligned with the orientation of the zebrafish (white dot in **Figure 2B**).

In control zebrafish, brain neural activity (red lines) was well synchronized with spinal cord activity, indicating a connected nervous system network where brain neural activation stimulates the spinal cord and motor neurons to fire together (**Figure 2C**). In contrast, *napb* CR exhibited neural hyperactivity in the brain region (**Figure 2D-F**). The number of zebrafish with brain region hyperactivity was significantly increased in the *napb* CR group (**Figure 2G**). Altogether, neural hyperactivity in the brain region of *napb* CR is correlated with seizure-like behavior and an epileptic phenotype (**Figure 1C-E**).

### MEA detects PTZ-induced seizure-like hyperexcitability

We further investigated whether MEA could detect the hyperexcitability of zebrafish induced by PTZ (**Figure 3A**). PTZ treatment significantly increased the weighted mean firing rate (**Figure 3B**), burst frequency (**Figure 3C**), and network burst frequency of neural network (**Figure 3D**) in the zebrafish control group, implying that PTZ-induced seizure-like hyperexcitability and epileptic hyperactivity can be effectively monitored using MEA.

**Figure 3.**
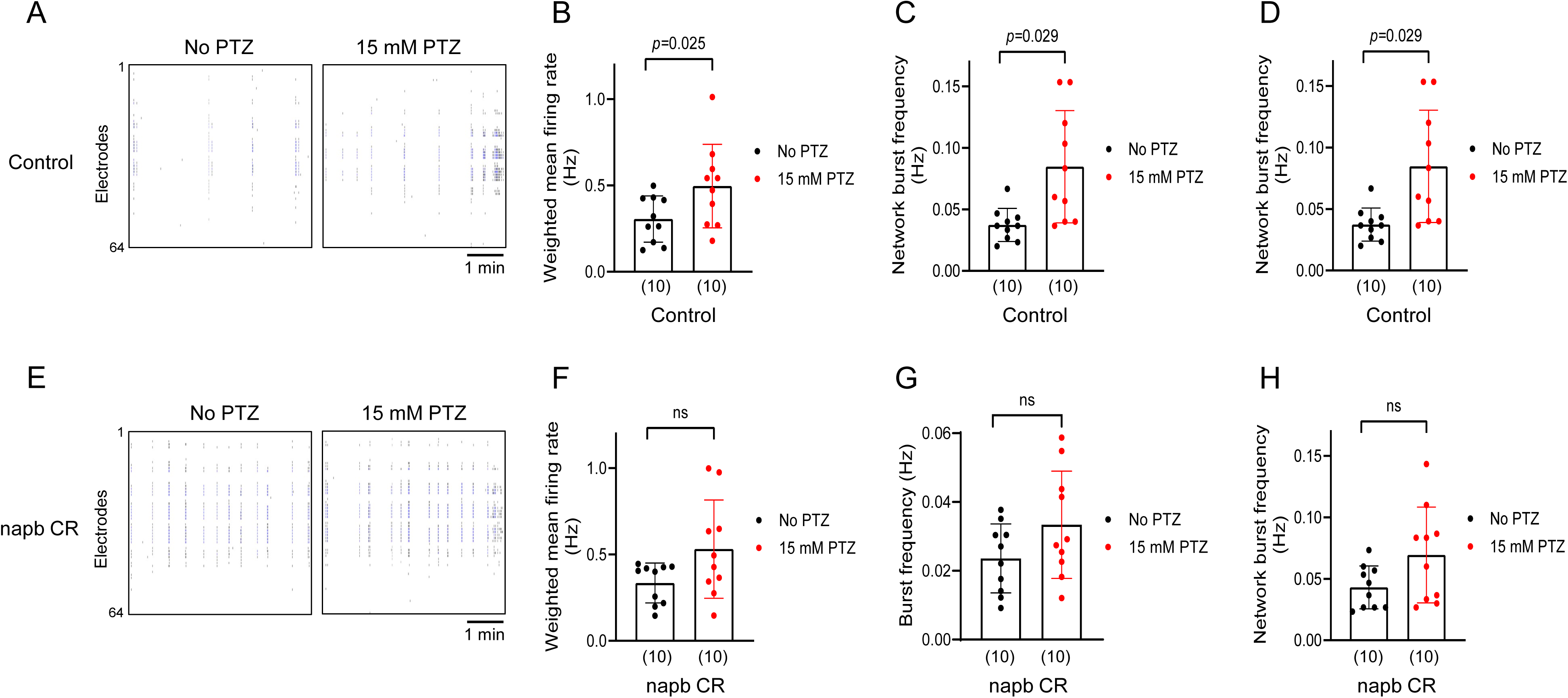
Seizure-like hyperexcitability induced by PTZ. Seizure-like neural hyperactivity was monitored using MEA. The neural activity of zebrafish larvae was recorded before and after treatment with 15 mM PTZ. (**A**) Raster plots of neural activity in control larvae before and after treatment with 15 mM PTZ. (**B-D**) Comparison of neural activity in zebrafish larvae in the absence or presence of 15 mM PTZ: control larvae (**B-D**) and *napb* CR (**E-H**). Quantification of (**B, F**) weighted mean firing rate (Hz), (**C,G**) burst frequency (Hz), and (**D, H**) network burst frequency of neural network (Hz). Data are presented as means ± SD for control larvae (**B-D**, n = 10) and *napb* CR (**F-H**, n = 10). The number of larvae tested is shown in parentheses. Repeated measure one-way ANOVA was used for statistical analysis to compare paired groups before and after PTZ treatment.

In *napb* CR (**Figure 3E-H**), the effect of PTZ on the weighted mean firing rate (**Figure 3F**) and network burst frequency (**Figure 3H**) was less pronounced compared to control zebrafish larvae. Additionally, the burst frequency was not significantly affected by PTZ in *napb* CR (**Figure 3G**). These findings indicate that PTZ may have a reduced ability to induce epileptic hyperactivity in NAPB zebrafish model, likely due to their already elevated hyperexcitability. This observation correlates with the seizure-like behavior phenotype observed in NAPB zebrafish model (**Figure 1F-H**).

## Discussion

Comorbidity is prevalent in ASD, including conditions such as attention-deficit hyperactivity disorder (ADHD) and epilepsy, both of which are linked to neuronal hyperexcitability^2^. Animal models for ASD-related epilepsy are essential for studying its pathophysiology. Our findings highlight the utility of *napb* zebrafish crispants as a model for studying ASD-associated epilepsy. Behavioral locomotor analyses showed significant hyperactivity, including increased swimming velocity and rotation frequency, consistent with seizure-like phenotypes. These results align with previous studies using zebrafish models of ASD-related epilepsy, where similar locomotor hyperactivity was observed, emphasizing that zebrafish are a robust system for modeling neural hyperexcitability and behavioral phenotypes associated with ASD and epilepsy^14,33^.

We applied MEA technology to monitor neural hyperexcitability in *napb* CR, integrating behavioral tests with electrophysiological functional assays to investigate the pathophysiology of epilepsy and associated neurological disorders. MEA recordings of *napb* CR zebrafish support previous findings of increased neural activity in zebrafish models of ASD. Notably, similar patterns of increased burst and network activity have been observed in MEA recordings of zebrafish epilepsy models^18,19^. These findings suggest that hyperexcitability may represent a shared neurophysiological hallmark across zebrafish models with comorbid epilepsy.

The PTZ response in *napb* CR is less sensitive, as *napb* CR already exhibit elevated hyperexcitability (**Figure 1**), highlighting their relevance in modeling the epileptic phenotype associated with *NAPB* mutations. The behavior of *napb* CR zebrafish in the presence of PTZ shows seizure-like behavior within the first 8 min of PTZ treatment; *napb* CR zebrafish have the significant higher distance, swimming velocity, and rotation frequency. However, the effect of PTZ diminished over time, potentially due to catatonic periods (**Supplementary Figure 6**) or it is likely that *napb* CR zebrafish are already in a hyperactive and seizure-like state, making additional hyperactivity induction by PTZ ineffective (**Figure 1C-E**). However, the underlying mechanism remains unclear and further investigations are required to elucidate the precise factors contributing to this observation. The use of F0 crispants allowed us to rapidly assess the functional impact of NAPB loss, significantly reducing experimental timelines compared to the generation of stable mutant lines. However, we recognize that F0 crispants may exhibit genetic mosaicism, which could influence phenotypic variability. Future studies utilizing stable mutant lines will be necessary to validate our findings and provide a more comprehensive understanding of NAPB function.

We acknowledge that MEA technology has some limitations in measuring spatial differences in neuronal activity due to uneven physical contact. Zebrafish larvae were laterally immobilized on the surface of the MEA electrodes using low-melting agarose (**Figure 2A**). While the MEA system enables simultaneous monitoring of neural activity from various locations across the zebrafish body, the three-dimensional shape of the larva introduces challenges in achieving uniform electrode contact. Specifically, the rounded structure of the head may prevent one or more electrodes in the head region from making optimal contact, potentially resulting in lower recorded activity in the corresponding heatmap. However, for our purposes, MEA is a highly effective and powerful tool for monitoring hyperexcitability and neural network formation in zebrafish. The observed synchronization between brain and spinal cord neural activity supports the interconnected and synchronized nervous system, where brain activation propagates to the spinal cord and motor neurons (**Supplementary Video 2**). The raster plot in **Figure 2C** highlights the synchronization of neural activity across the zebrafish body. However, not all 64 electrodes respond equally, as only those electrodes that are in close contact with the larva or sufficiently near to detect field potential changes contribute to the recorded signals. To address these observations comprehensively, we have included a supplementary video showcasing blinking signals and hyperexcitability in the brain area of the *napb* CR zebrafish (**Supplementary Video 2**). This visualization further supports the utility of MEA in detecting hyperactive neural activity in specific regions, even with the noted spatial limitations.

Epilepsy is a neurological disorder characterized by recurrent seizures^34^. Mutations in proteins involved in synapse and vesicle trafficking have been identified as contributing factors to the development of epilepsy^34,35^; for example, mutations in proteins such as synapsin, syntaxin 1B, and dynamin-1 are associated with epilepsy^34,35^. NAPB, selectively expressed in the brain, is a crucial cofactor of NSF ATPase for the disassembly of the SNARE complex^8^, thus playing a critical role in vesicle recycling and trafficking in neurons. However, *NAPB* deletion does not result in a dramatic phenotype in synaptic transmission at the cellular level^10^, and our studies have shown no defects in neuronal activity and calcium influx in *NAPB* mutant human stem cell-derived cortical neurons^4^, suggesting that cellular models may not adequately recapitulate the epileptic phenotype.

In zebrafish models of epilepsy, behavioral changes and hyperactivity can be quantified through various measures, such as total distance traveled, swimming velocity, and rotation frequency^11,15^. These parameters are reliable, functional indicators of seizure activity and neurological dysfunction. Zebrafish experiencing seizure-like hyperactivity typically exhibit increased total distance traveled and accelerated swimming velocity during seizure episodes^11,15^. Additionally, epileptic zebrafish often demonstrate an increased frequency of rotational movements, both clockwise and counterclockwise, which is a consequence of neural dysfunction and seizure severity^11,15^.

The established NAPB zebrafish model recapitulates key features of ASD-related epilepsy, including hyperactivity and neural hyperexcitability, and represents a promising animal model for investigating seizure-like phenotypes and epilepsy-related to *NAPB* mutations. These zebrafish crispants will be valuable for studying the pathophysiological mechanisms of ASD and epilepsy and screening potential therapeutic drugs. By combining behavioral tests with electrophysiological functional assays, the NAPB zebrafish model can provide comprehensive insights into the intervention strategies for epilepsy.

## Availability of data and material

The datasets supporting the conclusions of this article are available from the corresponding author on reasonable request.

## Ethics declarations Competing interests

The authors declare no conflict of interest.

## Ethics declaration and consent to participate

Zebrafish were maintained in standard conditions according to the Ministry of Public Health (MOPH), Qatar animal research guidelines and under an approved protocol by the Institutional Animal Care Committee (IACUC Office of Sidra Medicine, approval SIDRA-2023-001 and SIDRA-EXEMPT-2024-002). This study follows the ARRIVE Guidelines.

## Author Contributions

Conceptualization: K.C.S., W.H., G.A., L.W.S., S.I.D., Y.P.; Data curation: K.C.S., W.H.,, S.I.D.; Formal analysis: K.C.S., W.H.,, S.I.D., Y.P.; Investigation: K.C.S., W.H.,, S.I.D., Y.P.; Methodology: K.C.S., W.H.,, D.A., T.A., S.I.D., Y.P.; Resources: K.C.S., W.H., G.A., L.W.S.; Software: K.C.S., W.H., S.I.D.; Validation: K.C.S., W.H., S.I.D.;, Visualization: K.C.S., W.H., S.I.D., Y.P.; Project administration: L.W.S., S.I.D., Y.P.; Supervision: L.W.S., S.I.D., Y.P.; Funding acquisition: L.W.S., S.I.D., Y.P.; Writing - original draft: Y.P. and Writing - review & editing: K.C.S., W.H., G.A., L.W.S., S.I.D.

---

**Supplementary Figure 1.**
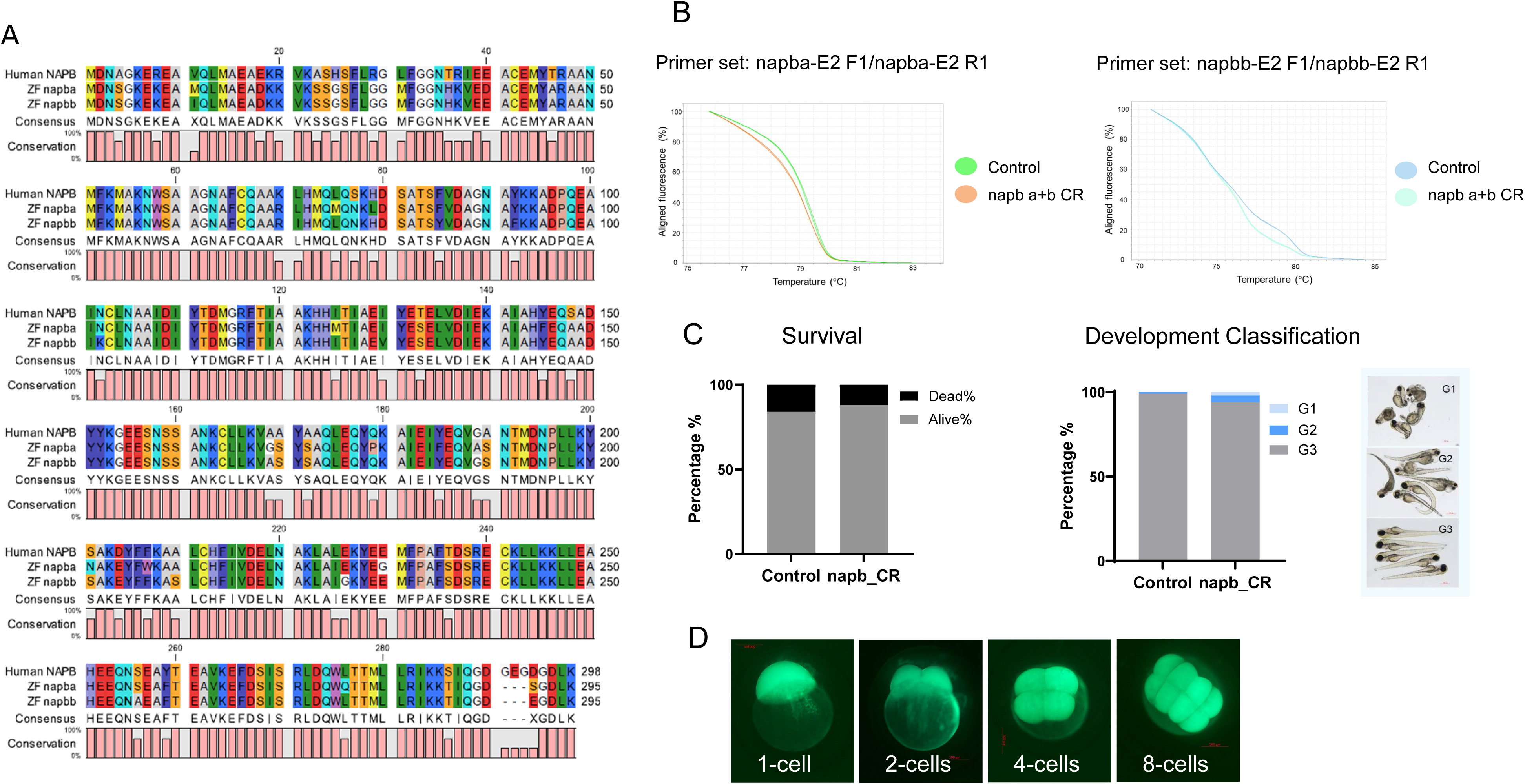

**Supplementary Figure 2.**
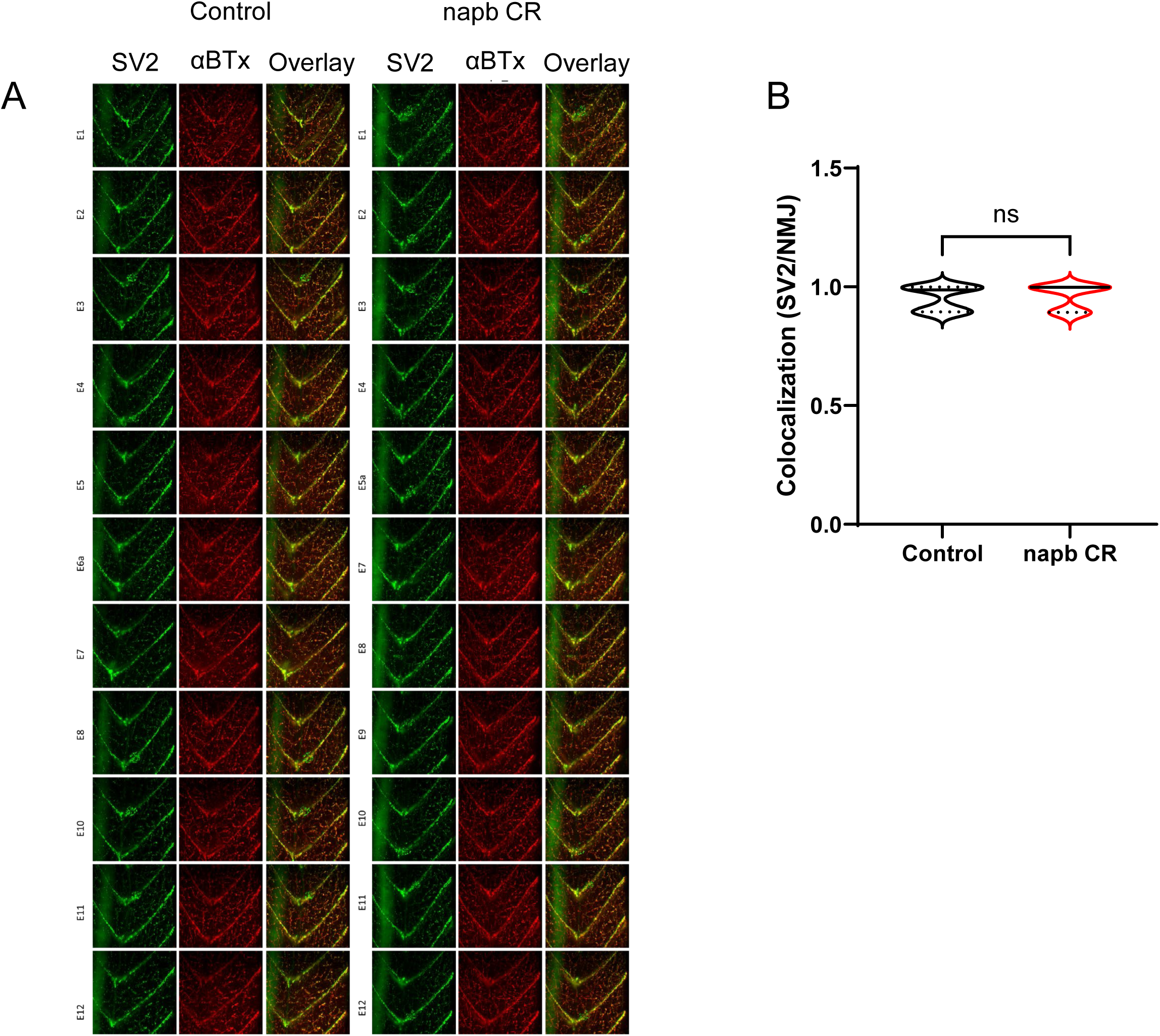

**Supplementary Figure 3.**
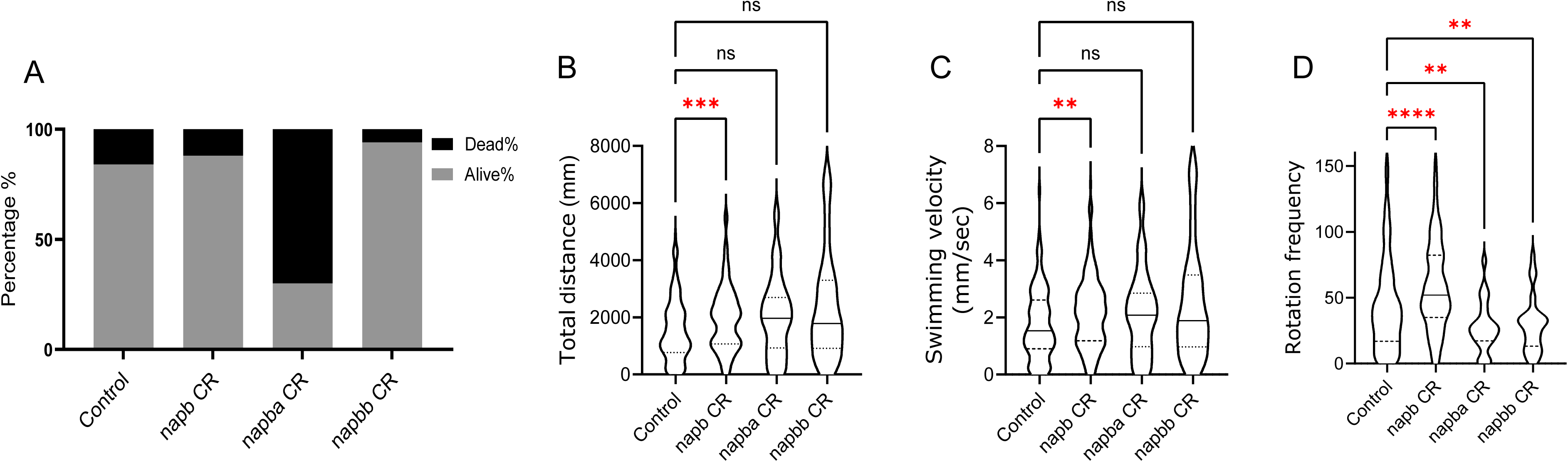

**Supplementary Figure 4.**
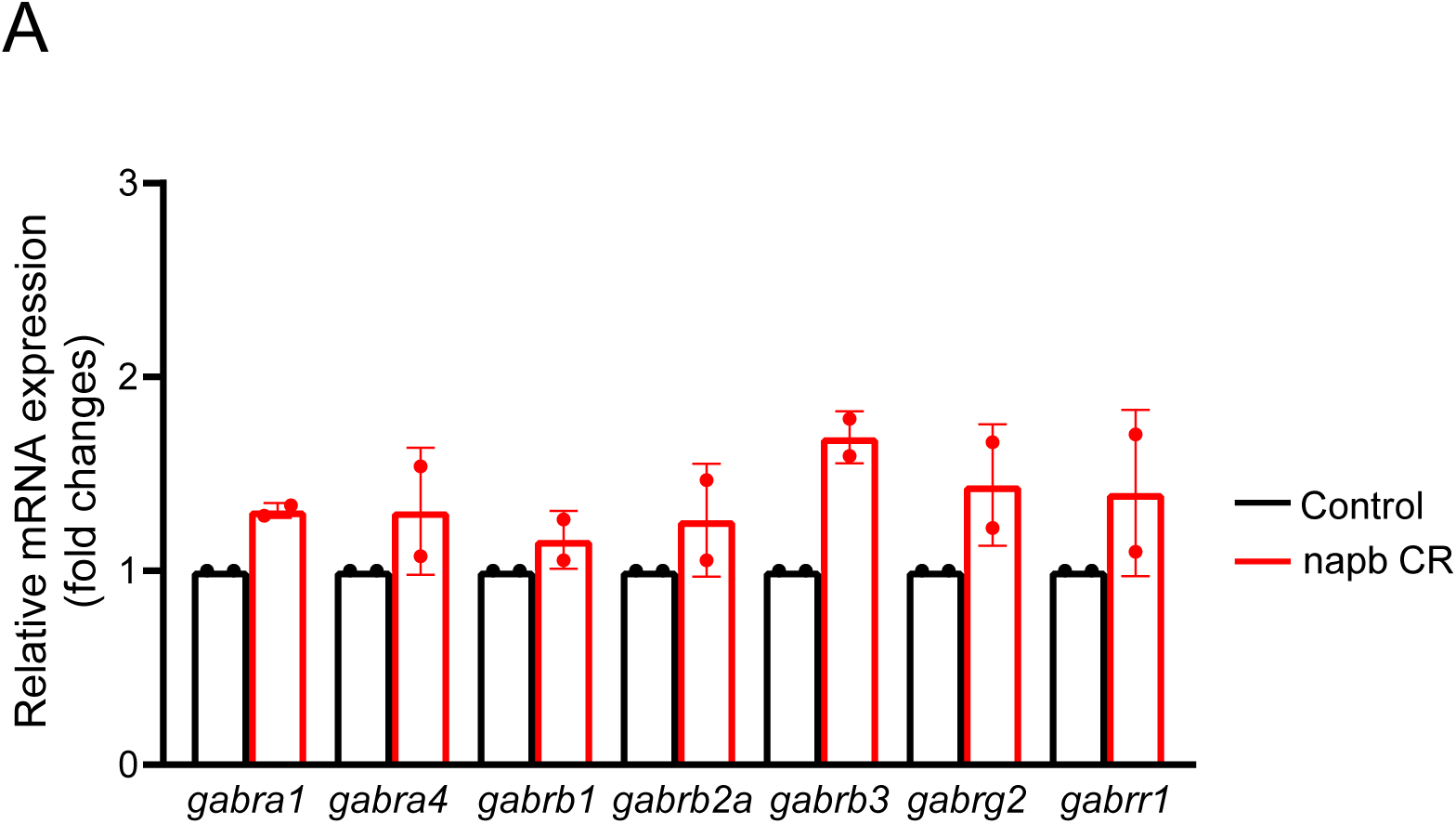

**Supplementary Figure 5.**
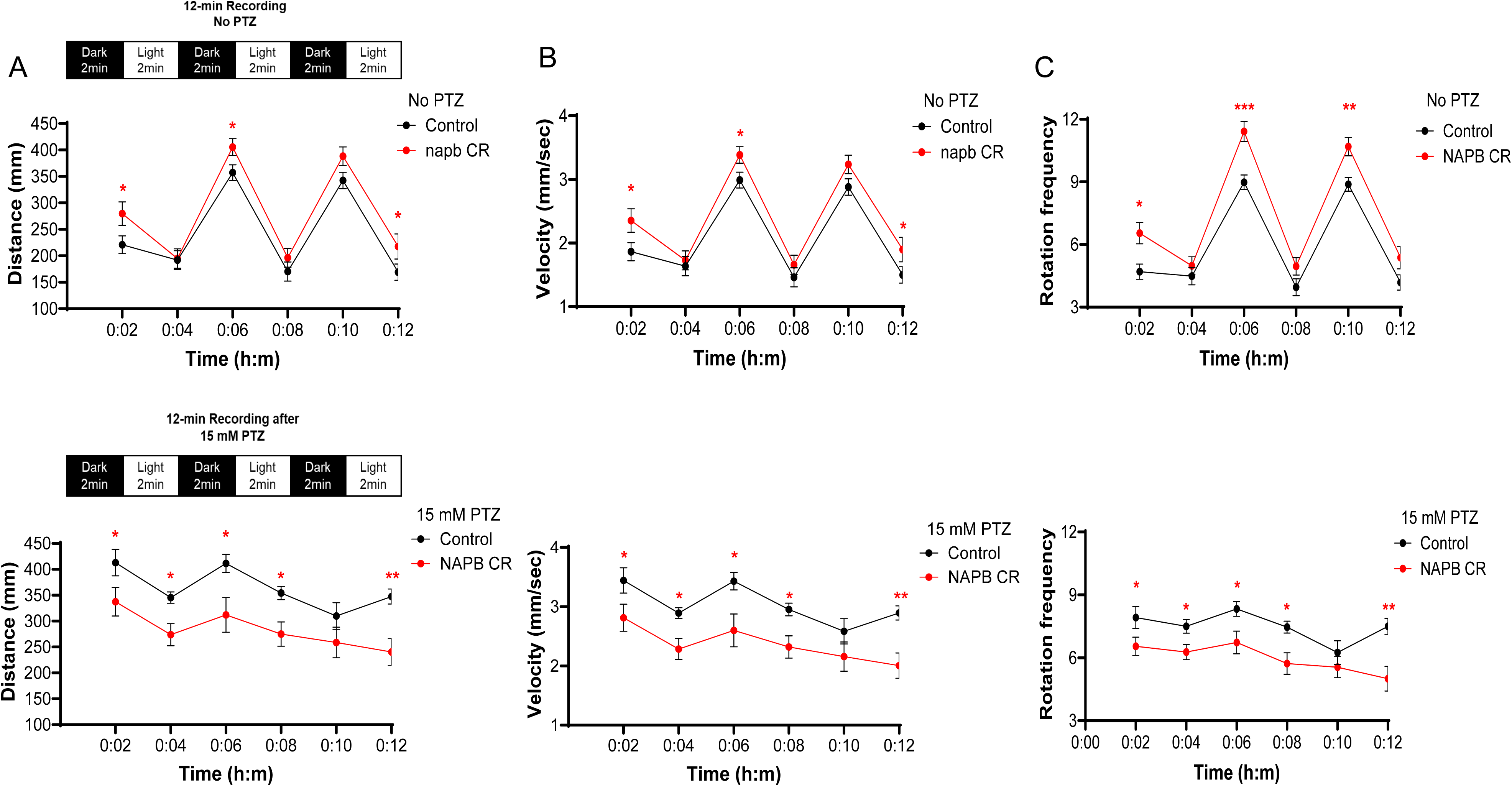

**Supplementary Figure 6.**
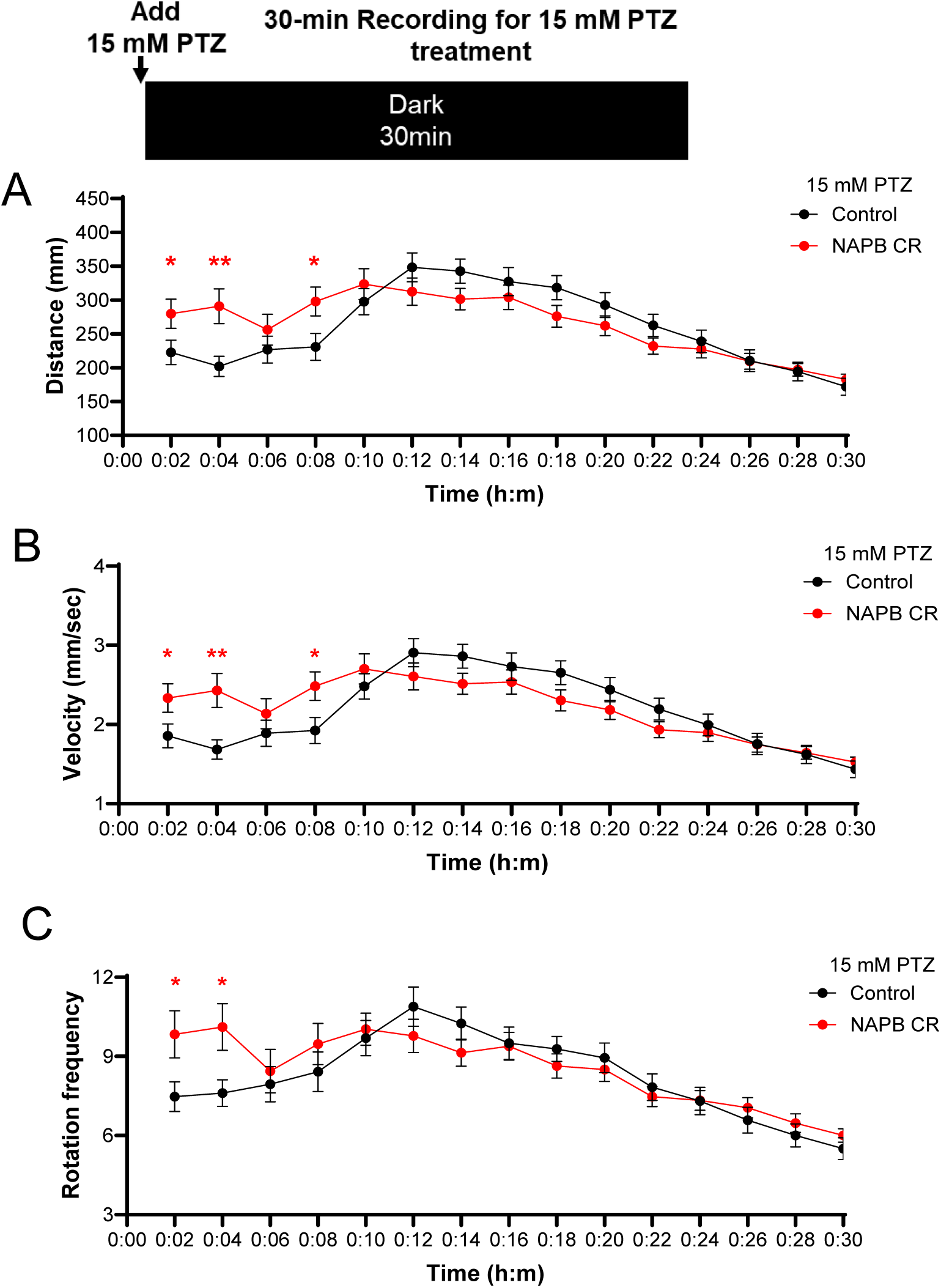

